# The neuroimaging correlates of depression established across six large-scale datasets

**DOI:** 10.1101/2025.07.02.660888

**Authors:** Kassandra Miyoko Hamilton, Xiaoke Luo, Ty Easley, Fyzeen Ahmad, Thomas Guo, Setthanan Jarukasemkit, Hailey Modi, Samuel Naranjo Rincon, Cabria Shelton, Lyn Stahl, Zijian Wang, Yuling Zhu, Petra Lenzini, Deanna M. Barch, Kayla Hannon, Janine Bijsterbosch

## Abstract

Neuroimaging data offers noninvasive insights into the structural and functional organization of the brain and is therefore commonly used to study the neuroimaging correlates of depression. To date, a substantial body of literature has suggested reduced size of subcortical regions and abnormal functional connectivity in frontal and default mode networks linked to depression. However, recent meta analyses have failed to identify significant converging correlates of depression across the literature such that a conclusive mapping of the neuroimaging correlates of depression remains elusive. Here we leveraged 23,417 participants across six datasets to comprehensively establish the neuroimaging correlates of depression. We found reductions in gray matter volume/ cortical surface area associated with depression in the frontal cortex, anterior cingulate, and insula, confirming prior studies showing the importance of prefrontal and default mode regions in depression. Our findings demonstrate multiple surprising results, including a lack of depression correlates in subcortical brain regions, significant depression correlates in somatomotor and visual regions, and limited functional connectivity findings. Overall, these results shed new light on key brain regions involved in the pathophysiology of depression, updating our understanding of the neuroimaging correlates of depression. We anticipate that these insights will inform further research into the role of sensorimotor and visual regions in depression and into the impact of heterogeneity on functional connectivity correlates of depression.

## Introduction

Over 11,000 papers have been published to date to investigate the structural and functional neuroimaging correlates of depression^1^. However, recent meta analyses attempting to collate results across prior studies have reported null findings^1–4^, indicating a troubling lack of consistency in neuroimaging correlates of depression across the substantive existing literature. Sampling variability arising from underpowered study samples that comprise each meta-analysis may explain these null findings^5^, and such within-cohort sample size challenges are often present even in large-scale studies of the neural correlates of depression^6,7^. As such, a conclusive mapping of the neuroimaging correlates of depression remains unknown. In this study, we aimed to comprehensively identify the most reliable structural and resting state functional neuroimaging correlates of depression across six large-scale datasets.

Beyond sampling variability in underpowered studies, there are several other challenges that may contribute to the observed inconsistencies in the literature on neuroimaging correlates of depression. For example, there are multiple self-reported phenotypic definitions of depression in common use, which are broadly based on either symptom severity or trait-level predisposition indexed using the neuroticism personality trait^8,9^. Furthermore, the literature on neuroimaging correlates of depression has largely focused on higher-order cortical systems^10^ and subcortical regions^11^, which may have systematically resulted in under-reporting in other regions of the brain. To address these challenges, we determined the unbiased whole-brain spatial distribution of the structural and functional neuroimaging correlates reliably associated with symptom severity versus predisposition phenotypes of depression.

We used six state-of-the-art large-scale datasets to comprehensively map the parcellated structural and resting state functional neuroimaging correlates of depression. Each dataset was analyzed separately and estimates were subsequently meta-analytically combined to establish the most reliable neuroimaging correlates of depression robust to sampling variability, phenotypic definitions, and regional bias (Fig. 1). We purposefully chose to treat the six datasets included in this study separately rather than harmonizing all data into one mega sample. This approach was selected because perfect harmonization is not feasible^12^, especially when factors of importance (such as age range) are fully co-linear with dataset separation. Furthermore, non-removal of study differences (e.g., in relation to pre-processing, sample definition, depression instruments, etc.) offers a testbed for the type of realistic study-to-study variation observed in the literature.

**Figure 1.**
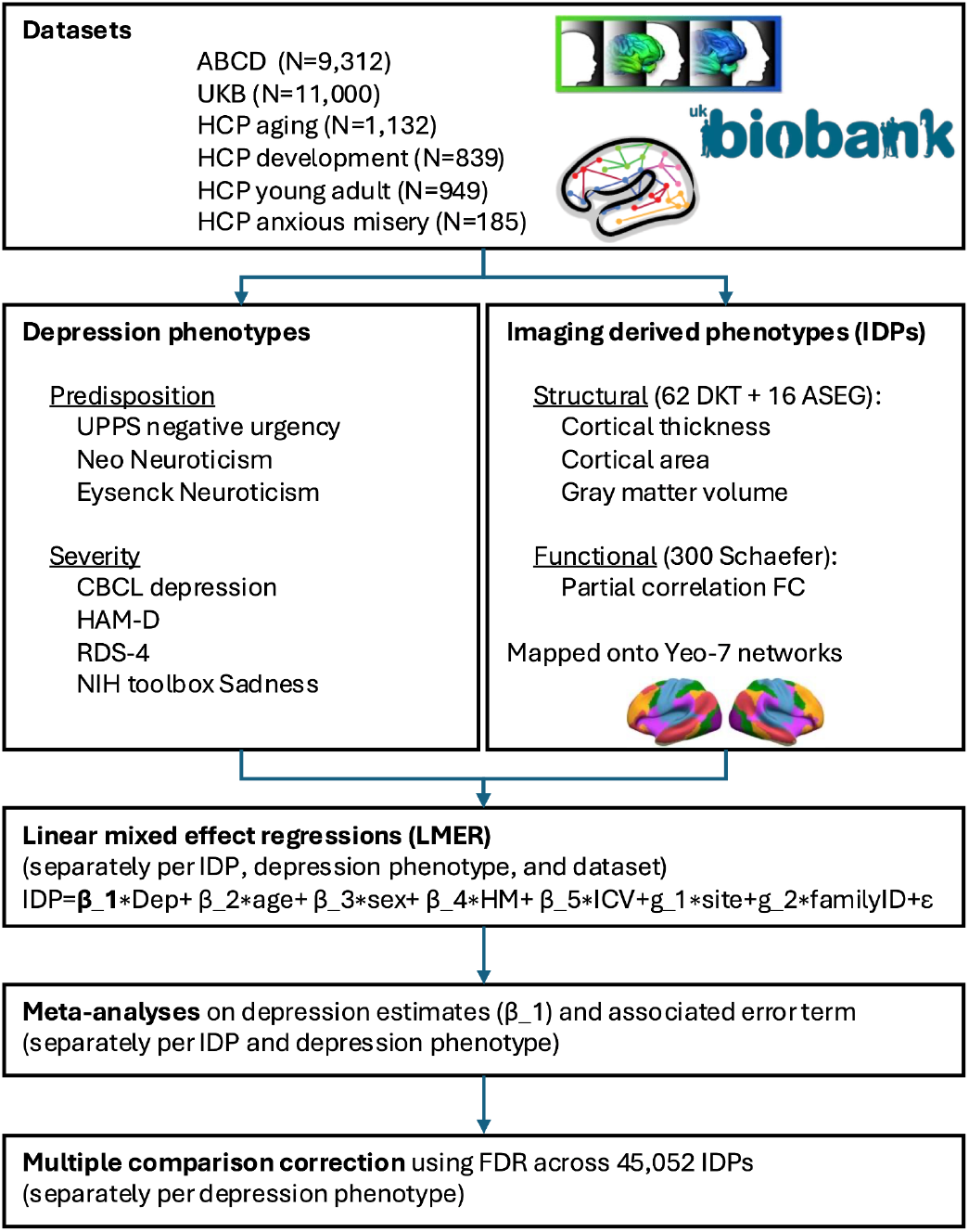
Overview of main analysis steps. From each of the 6 datasets, we extract one available depression severity phenotype and one available depression predisposition phenotype, and 45,052 matched imaging derived phenotypes. To estimate the association of each IDP with each depression phenotype, univariate within-dataset analyses were performed using linear mixed effect regressions and the results were entered into univariate meta-analyses to combine across datasets, followed by multiple comparison correction using false discovery rate (FDR). See the Online Methods for more information.

Our findings identified significant structural neuroimaging correlates of depression. Brain regions with the strongest and most robust reductions in gray matter volume/ cortical surface area associated with depression included the frontal cortex, anterior cingulate and insula. Importantly, our findings did not indicate any significant resting state functional neuroimaging correlates of depression severity, casting doubt on the mechanistic theory of depression as a dysconnectivity syndrome. With regards to the whole-brain spatial distribution, significant structural neuroimaging correlates of depression included visual and somatomotor regions (e.g., reduced volume/area in paracentral, postcentral, precentral, fusiform, and pericalcarine regions), but no subcortical regions, shedding new light on key brain regions involved in the pathophysiology of depression. Taken together, this study substantially updates our understanding of the neuroimaging correlates of depression.

### Robust structural neuroimaging correlates of depression

To establish the structural and functional neuroimaging correlates of depression, we comprehensively analyzed six existing datasets including the Adolescent Brain Cognitive Development (ABCD) study^13^, The UK Biobank (UKB) study^14^, the Human Connectome Project (HCP) Young Adult study^15^, the HCP Development study^16^, the HCP Aging study^17^, and the HCP Dimensional Connectomics of Anxious Misery study^18^. Depression severity was quantified based on available metrics in each dataset, using either the Hamilton Depression Rating Scale^19^ (ANXPE), NIH Toolbox Sadness^20^ (HCP-YA, HCP-A), Child Behavior Checklist (CBCL) Depression subscale^21^ (HCP-D, ABCD), or Recent Depressive Symptoms (RDS) scale^22^ (UKB). Depression predisposition was similarly quantified using dataset-specific measures, namely the Eysenck Neuroticism^23^ (UKB), NEO five-factors inventory questionnaire Neuroticism subscale (HCP-YA, HCP-A, ANXPE), or UPPS negative urgency^24^ (HCP-D and ABCD). Parcellated structural MRI metrics (gray matter volume, cortical thickness, cortical surface area) were calculated for 62 brain regions in the Desikan-Killiany-Tourville (DKT)^25^ atlas, and - for gray matter volume - 16 subcortical regions in the ASEG^26^ atlas. Functional connectivity was defined as the partial correlation between pairs of parcel timeseries using the Schaefer^27^ 300-dimensional atlas. Mass univariate linear mixed-effects regression (LMER) models were fit to separately estimate each imaging derived phenotype based on one of the depression phenotypes (severity or predisposition) and confounds (age, sex, total intracranial volume, head motion, imaging site, and family group, as relevant for each dataset). LMER effect size and error estimates from each of the six datasets were subsequently entered into mass univariate meta analyses to calculate overall brain-depression effect sizes separately for each imaging derived phenotype and each depression phenotype. False discovery rate (FDR) correction was performed to control meta analysis results for multiple comparisons across all structural and functional imaging measures within each depression phenotype, such that no significance is reported at the study level.

The results revealed robust structural neuroimaging correlates of depression. The most consistent neural correlates of depression were reduced gray matter volume/ cortical surface area in the superior frontal cortex, middle frontal cortex, orbitofrontal cortex, rostral anterior cingulate and insula (Fig. 2; full set of results included in the supplementary information). Notably, among significant regional findings, there was substantial agreement both bilaterally and between gray matter volume and cortical surface area (Table 1). However, no significant meta analytical results were observed for cortical thickness, which may be explained by recent findings revealing extensive heterogeneity in cortical thickness changes associated with depression^28^. Overall, these findings are consistent with prior voxel-based meta analytical studies^29–32^, and point to the importance of frontal regions in depression.

**Table 1:** Overview of significant structural meta analytical effect sizes. Results that pass FDR-corrected p<0.05 are highlighted in blue font. Counts of significant entries are summarized per row (# Sig) and per column.

| Region | Yeo Network | Severity |  |  |  | Predisposition |  |  |  | # Sig |
| --- | --- | --- | --- | --- | --- | --- | --- | --- | --- | --- |
|  |  | Surface Area |  | Volume |  | Surface Area |  | Volume |  |  |
|  |  | Left | Right | Left | Right | Left | Right | Left | Right |  |
| Superior frontal | Default | -0.0584 | -0.0662 | -0.0517 | -0.0577 | -0.0584 | -0.0561 | -0.0428 | -0.0438 | 7 |
| Lateral orbitofrontal | Limbic | -0.0400 | -0.0421 | -0.0363 | -0.0373 | -0.0481 | -0.0464 | -0.0371 | -0.0374 | 6 |
| Rostral middle frontal | Frontoparietal | -0.0500 | -0.0459 | -0.0384 | -0.0399 | -0.0485 | -0.0489 | -0.0412 | -0.0398 | 6 |
| Rostral anterior cingulate | Default | -0.0367 | -0.0249 | -0.0341 | -0.0169 | -0.0367 | -0.0382 | -0.0398 | -0.0305 | 5 |
| Caudal middle frontal | Frontoparietal | -0.0210 | -0.0289 | -0.0067 | -0.0286 | -0.0404 | -0.0382 | -0.0359 | -0.0355 | 4 |
| Inferior parietal | Default | -0.0503 | -0.0495 | -0.0472 | -0.0461 | -0.0331 | -0.0179 | -0.0303 | -0.0204 | 4 |
| Medial orbitofrontal | Limbic | -0.0162 | -0.0355 | 0.0058 | -0.0208 | -0.0381 | -0.0404 | -0.0357 | -0.0261 | 4 |
| Precuneus | Default | -0.0481 | -0.0478 | -0.0493 | -0.0476 | -0.0301 | -0.0311 | -0.0288 | -0.0181 | 4 |
| Middle temporal | Default | -0.0438 | -0.0544 | -0.0258 | -0.0489 | -0.0205 | -0.0351 | -0.0213 | -0.0262 | 3 |
| Paracentral | Somatomotor | -0.0427 | -0.0263 | -0.0453 | -0.0341 | -0.0158 | -0.0123 | -0.0217 | -0.0084 | 3 |
| Supramarginal | Ventral Attention | -0.0443 | -0.0378 | -0.0422 | -0.0394 | -0.0264 | -0.0461 | -0.0228 | -0.0344 | 3 |
| Caudal anterior cingulate | Ventral Attention | -0.0383 | -0.0166 | -0.0315 | -0.0150 | -0.0324 | -0.0204 | -0.0302 | -0.0150 | 2 |
| Insula | Ventral Attention | -0.0437 | -0.0325 | -0.0408 | -0.0366 | -0.0291 | -0.0304 | -0.0286 | -0.0297 | 2 |
| Pericalcarine | Visual | -0.0183 | -0.0088 | -0.0131 | -0.0070 | -0.0311 | -0.0202 | -0.0310 | -0.0177 | 2 |
| Postcentral | Somatomotor | -0.0296 | -0.0494 | -0.0198 | -0.0436 | -0.0231 | -0.0400 | -0.0031 | -0.0267 | 2 |
| Fusiform | Visual | -0.0338 | -0.0393 | -0.0308 | -0.0331 | -0.0326 | -0.0321 | -0.0313 | -0.0244 | 1 |
| Posterior cingulate | Ventral Attention | -0.0328 | -0.0264 | -0.0312 | -0.0209 | -0.0077 | -0.0210 | -0.0131 | -0.0099 | 1 |
| Precentral | Somatomotor | -0.0239 | -0.0407 | -0.0328 | -0.0465 | -0.0257 | -0.0295 | -0.0171 | -0.0220 | 1 |
| # significant |  | 10 | 8 | 9 | 11 | 6 | 6 | 6 | 4 |  |
| # significant |  | 38 |  |  |  | 22 |  |  |  | 60 |

**Figure 2.**
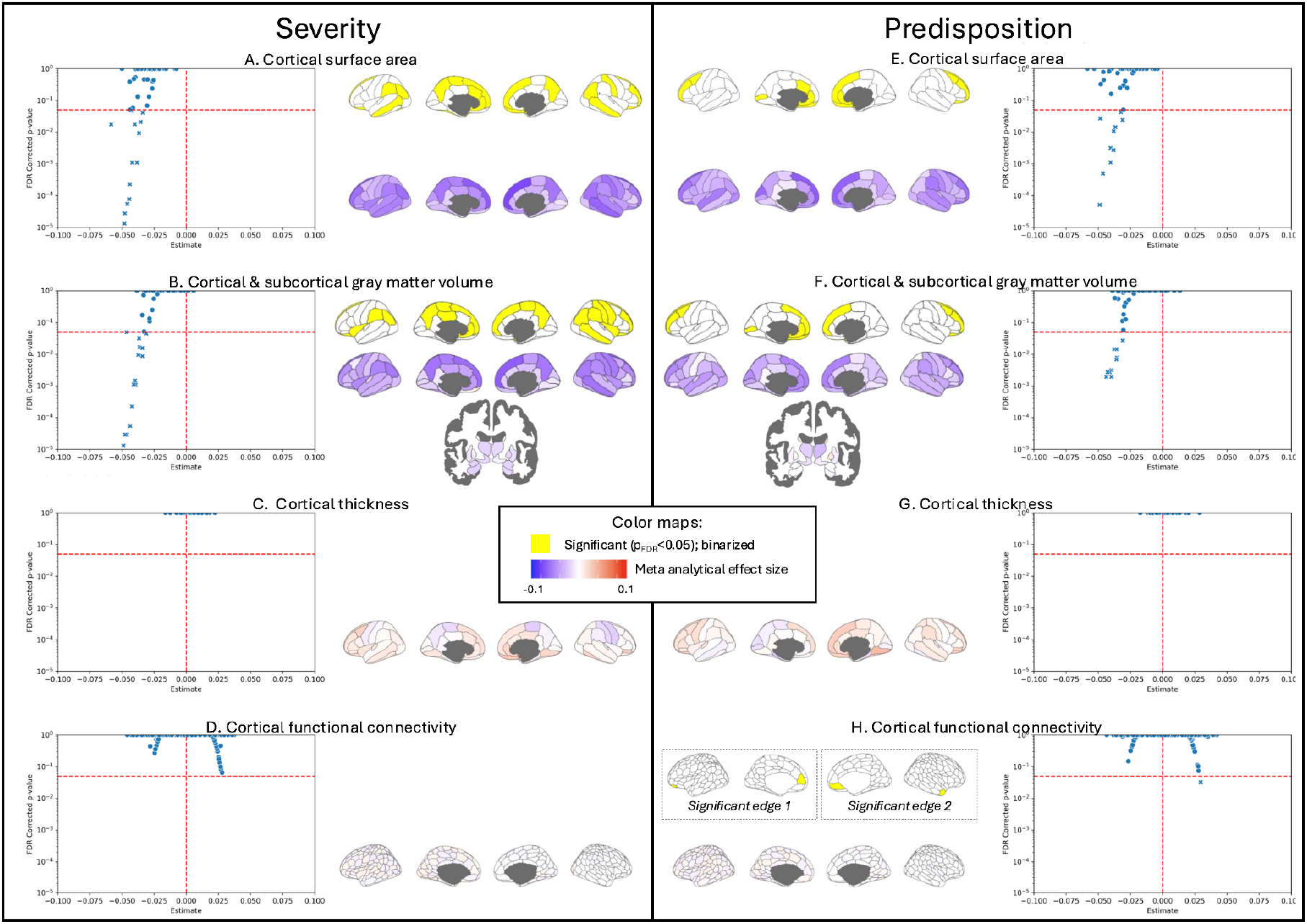
Neuroimaging correlates of depression collated across six large-scale datasets for (A-D) depression severity and (E-H) predisposition phenotypes. Volcano plots reveal meta analytical estimates, with significant imaging measures indicated by crosses. Results are shown on the parcellated cortical surface unthresholded (cold-warm colorbar) and after thresholding (yellow; where significant). For functional connectivity, cortical surface figures show the absolute average across a row of the full correlation matrix with the sign determined by the mode of the sign across the row (e.g., negative sign if more edges of a brain region are negative than positive).

Importantly, the meta analysis did not indicate significant gray matter volume changes associated with depression severity or predisposition in any of the 16 subcortical regions. Notably, none of the subcortical results achieved significance prior to multiple comparison correction either, although uncorrected trends were observed in the bilateral amygdala, left hippocampus, and right accumbens (p_uncorrected_=0.07; see supplementary information). It is possible that this lack of subcortical associations may be driven by a nonlinear relationship between subcortical volume and depression severity/predisposition that is obscured with the linear statistics adopted here. Such a nonlinear association may be more likely when using population datasets compared to traditional patient-control studies of depression, and therefore warrant future investigation.

### No robust functional correlates of depression

Surprisingly, our meta-analysis found no functional connectivity results that reached significance after correction for multiple comparisons for depression severity (Fig. 2D), and only two functional connectivity edges that reached significance for depression predisposition (Fig. 2H). Specifically, depression predisposition was associated with increased functional connectivity between the right anterior temporal cortex and the right medial prefrontal cortex and with increased functional connectivity between the left medial prefrontal cortex and the left ventral prefrontal cortex. These findings are consistent with prior work linking depression to hyperconnectivity within the default mode network and between the default mode network and frontoparietal network^10^. Prior work has furthermore linked increased activity within the default mode network with increased rumination^33,34^ and self-referential thinking^35^. As such, our limited functional connectivity findings support the role of rumination in depression predisposition^34^.

Overall, the lack of substantive functional connectivity findings was surprising given the extensive prior literature highlighting functional connectivity alterations as a key descriptor of depression etiology^10^, which has led to major depressive disorder being described as a dysconnectivity syndrome^36^. Notably, the multiple comparison burden was combined across all structural and functional imaging measures and therefore cannot explain this difference. It is possible that the functional correlates of depression may be more heterogeneous across patients than structural correlates^37–39^. These results call into question the description of depression in light of dysconnectivity mechanisms, and raise the importance of robust studies of depression heterogeneity focused on functional connectivity.

To further enable quantification of direct comparisons between the effect sizes of structural and functional neuroimaging correlates of depression, we performed a two-way ANOVA with a main effect for imaging metric type (4 levels; gray matter volume, cortical surface area, cortical thickness, functional connectivity), a main effect for depression phenotype (2 levels; severity, predisposition), and the interaction effect (imaging metric type x depression phenotype) on the absolute values of the meta-analytical effect sizes as the inputs. Importantly, this analysis was performed on all effect sizes (regardless of significance after multiple comparison correction) and can therefore pick up broader trends beyond the univariate results described above.

The results revealed a significant main effect of imaging metric type (F=1700.3, p=0), a significant main effect of depression phenotype (F=15.7, p=7.42*10^−5^), and a significant interaction effect (F=12.6, p=3.02*10^−8^). Post-hoc results confirmed significantly higher effect sizes for depression severity compared to predisposition (especially for gray matter volume and cortical surface area). Post-hoc results further indicated significant differences between each pair of imaging metric types, with especially larger effect sizes for gray matter volume and cortical surface area as compared to functional connectivity and cortical thickness (Fig. 3).

**Figure 3.**
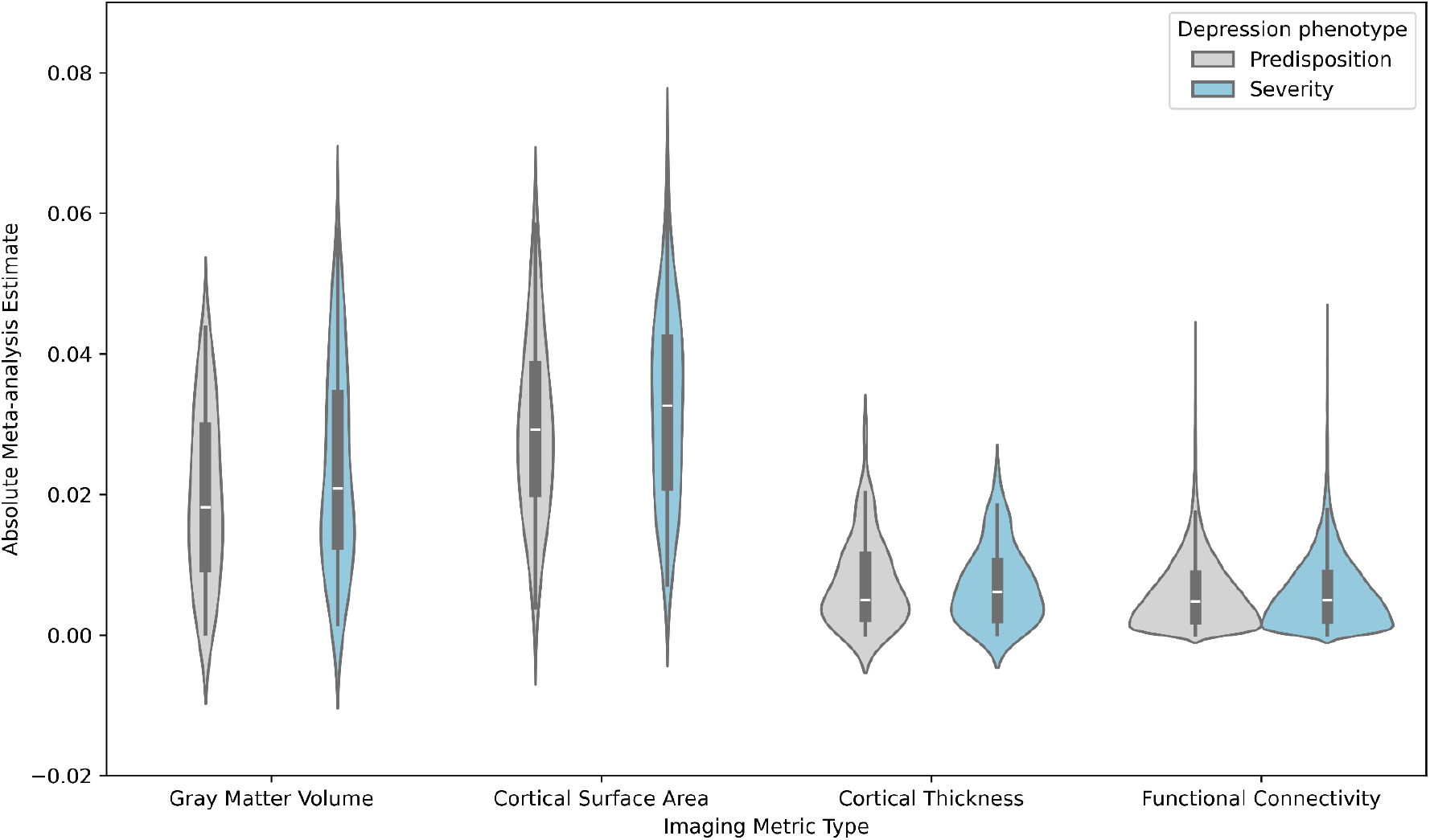
Meta analytical estimates (absolute) as a function of imaging metric type (x-axis) and depression phenotype. Results show larger effects for severity (blue) compared to predisposition (gray), especially for gray matter volume and surface area.

### Visual and somatomotor involvement in depression

The literature on the structural and functional neuroimaging correlates of depression has largely focused on subcortical regions (e.g., amygdala, hippocampus) and higher order cortical networks (e.g., regions of the default mode, fronto-parietal, and executive control networks)^10,11^. We aimed to perform an unbiased search for the structural and functional neuroimaging correlates of depression across the whole brain. Qualitatively, the meta analytical results summarized in Fig. 2 reveal a diffuse whole-brain pattern of associations with depression^40^. To systematically summarize the spatial distribution of neuroimaging correlates of depression, we assigned each parcel (from DKT or Schaefer atlases) to one of 7 cortical networks as defined by Yeo et al^41^ or to an eighth combined subcortical network (ASEG) based on maximal spatial overlap. We then performed ANOVAs with a main effect for Yeo network (7 or 8 levels; depending on the inclusion/exclusion of subcortical regions), a main effect for depression phenotype (2 levels; severity and predisposition), and the interaction effect (Yeo network x depression phenotype) on the absolute values of the meta-analytical effect sizes as the inputs. Analyses were performed separately for each of the two imaging metric types with significant meta analytical results (namely, gray matter volume and cortical surface area).

The results revealed a significant main effect of Yeo network for gray matter volume (F=10.1, p=3.5*10^−10^), which was driven by relatively lower effect sizes for subcortical regions (Fig. 4A). This finding is surprising given prior research indicating associations between depression and volume of several subcortical regions^6^ including the hippocampus^42^. Nevertheless, our findings are consistent with recent work showing chance-level multivariate classification of major depressive disorder, where subcortical volumes became uninformative for classification after careful harmonization for site effects^43^. It is possible that the wide variation of age ranges across the datasets may contribute to this surprising observation, given that hippocampal volume loss has been linked to cumulative depressive burden across lifetime episodes^44,45^ and may be absent in first episode patients^6^. Alternatively, future work may wish to assess nonlinear associations between subcortical (e.g., hippocampal) volume and depression in population data.

**Figure 4.**
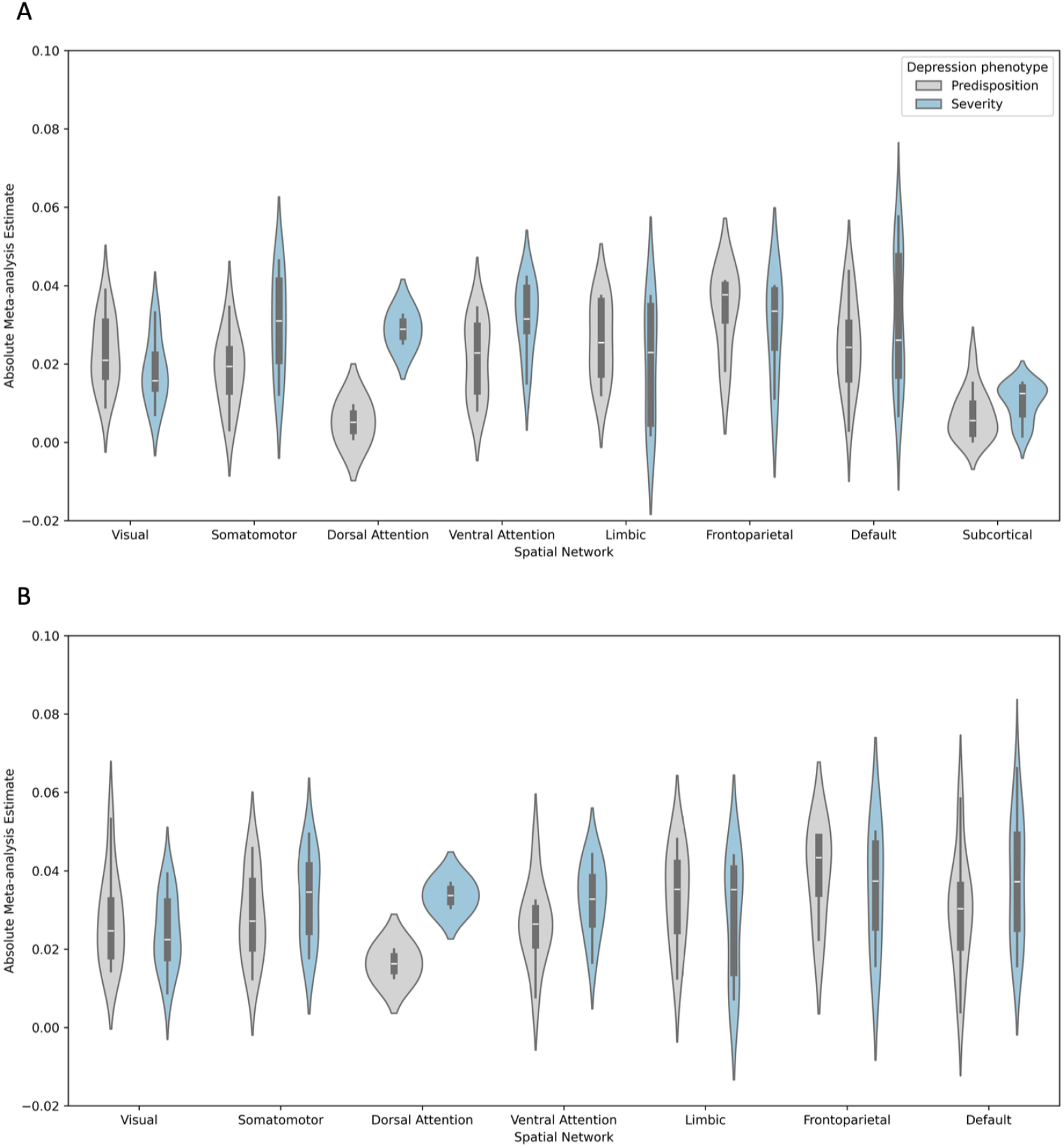
Neuroimaging correlates of depression as a function of spatial network organization (x-axis categories) and depression phenotype (gray = predisposition, blue = severity), shown separately for (A) gray matter volume, (B) cortical surface area. The white dots in each violin plot represent the median and the thick gray bar in the center represents the interquartile range. On each side of the gray line is a kernel density estimation to show the distribution shape of the data. Notably, the ‘limbic’ network in figure 4 refers to the Yeo-7 cortical definition of limbic regions (i.e., medial and lateral orbitofrontal cortex, inferior temporal cortex, and entorhinal cortex) and does not include subcortical contributions.

Interestingly, post-hoc comparisons of the ANOVA results for gray matter volume and cortical surface area did not indicate any significant differences between cortical networks (p>0.25). Although more brain regions in the default mode, frontoparietal, and limbic networks showed significant associations with depression (Table 1), multiple robust significant structural associations with depression were also observed for regions in the somatomotor and visual networks. Specifically, depression severity was associated with reduced gray matter volume and/or cortical surface area in paracentral, postcentral, precentral, and fusiform regions, whereas depression predisposition was associated with reduced gray matter volume and/or cortical surface area in the pericalcarine region. Recent studies have started to acknowledge and discuss the role of the visual and somatomotor networks in depression^46–49^, but attention on these findings remains limited. Our results substantiate structural associations of the visual and somatomotor networks in depression, highlighting the need to improve unbiased reporting and future research into potential explanatory mechanisms. In relation to the role of somatomotor networks in depression, recent studies have revealed the transdiagnostic nature of somatomotor abnormalities across multiple psychiatric and cognitive domains^50–53^. Taken together, there is a need for increased research focus on the role of the visual and somatomotor regions in depression.

### Unique neuroimaging correlates of severity versus predisposition

Variability in depression phenotypes is one potential reason for the observed inconsistencies across the literature on neuroimaging correlates of depression. We directly compared the neuroimaging correlates across two classes of depression phenotypes, namely self-reported instruments that measure depression severity versus self-reported instruments that measure the personality trait neuroticism as a proxy for depression predisposition. Notably, the exact instrument within each class of depression phenotype varied across the datasets (see Online Methods for details). To assess systematic differences in meta analytic effect sizes, the same ANOVAs for gray matter volume and cortical surface area described above - which included a main effect for depression phenotype and an interaction effect (depression phenotype by network) - were used.

In the ANOVAs for gray matter volume, both the main effect of depression phenotype (F=5.16, p=2.46*10^−2^) and the interaction effect (depression phenotype by network; F=2.19, p=3.89*10^−2^) were significant. Post hoc comparisons indicated that the main effect was driven by larger effect sizes for severity compared to predisposition. This is consistent with the univariate thresholded results, which indicated 38 significant hits for severity as compared to 22 significant hits for predisposition (Table 1). Interaction effects were driven by subcortical regions as discussed above. For cortical surface area neither the main effect of depression phenotype (F=1.83, p=0.17) nor the interaction effect (F=1.09, p=0.37) reached significance.

We note that potentially higher effect sizes may be expected when using clinician-rated diagnostic tools such as the Hamilton Depression Rating scale (HAM-D) or the structured clinical interview for diagnosis (SCID)^54–57^, even compared to self-reported indices of symptom severity. Although we did leverage clinician-rated instruments where available (in HCP-ANXPE), there is currently a lack of sufficiently large-scale neuroimaging cohorts with extensive clinician-rated phenotyping. Beyond clinician-rated severity scores, phenotypic variability in symptom type forms another source of potential bias in studies of the brain basis of depression. Future work may wish to explore the meta analytical brain basis of specific symptom domains.

## Conclusion

In this study, we comprehensively analyzed the structural and functional neuroimaging correlates of depression across six large-scale datasets and meta-analytically combined results to establish the most reliable brain regions involved in the psychophysiology of depression. Our findings revealed that depression was associated with significant reductions in gray matter volume and cortical surface area in superior frontal cortex, middle frontal cortex, orbitofrontal cortex, rostral anterior cingulate and insula. Surprisingly, we did not observe any evidence for depression correlates in subcortical brain areas, and very limited evidence for depression correlates in functional connectivity. Although the majority of findings involved default mode, frontoparietal, and limbic regions, multiple unexpected structural associations with depression were also observed in somatomotor (paracentral, postcentral, and precentral) and visual (fusiform and pericalcarine) regions. Taken together, this study is the first to robustly map the neuroimaging correlates of depression across well powered studies. Importantly, these findings substantially update our understanding of the neuroimaging correlates of depression by highlighting the importance of structural rather than functional associations and uncovering key somatomotor and visual contributions.

## Data sharing statement

All datasets used in this manuscript have been shared online:

- UKB data are available following an access application process: https://www.ukbiobank.ac.uk/enable-your-research/apply-for-access. This research was performed under UK Biobank application number 47267.
- ABCD data was available from the NIMH Data Archive when data were obtained for this study.
- HCP-A, HCP-D, ABCD and ANXPE (referred to as Dimensional Connectomics of Anxious Misery [DCAM]) data are available from the NIMH Data Archive collectively under the header of ‘CCF Data from the Human Connectome Projects’.
- HCP-YA data are available from the connectomeDB: https://db.humanconnectome.org/.

## Code sharing statement

All code used in this paper are available on Github:

## Author contributions

Conceptualization: KH, KMH, JB

Data Curation: XL, KMH

Formal Analysis: KMH, XL, TE, FA, TG, SJ, HM, SNR, CS, LS, PW, YZ, PL

Funding Acquisition: KMH, JB

Methodology: TE, PL, LS, XL

Supervision: KH, JB

Visualization: XL, KMH, SJ

Writing – Original Draft: KH, JB, KMH, XL

Writing – Review & Editing: KMH, XL, TE, FA, TG, SJ, HM, SNR, CS, LS, PW, YZ, PL, KH, JB

## Footnotes

1 Based on a PubMed search for the keyword string ‘neuroimaging depression’.

## References

1. Saberi, A., Mohammadi, E., Zarei, M., Eickhoff, S. B. & Tahmasian, M. Structural and functional neuroimaging of late-life depression: a coordinate-based meta-analysis. Brain Imaging Behav. (2021) doi:10.1007/s11682-021-00494-9.

2. Winter, N. R. et al. Quantifying Deviations of Brain Structure and Function in Major Depressive Disorder Across Neuroimaging Modalities. JAMA Psychiatry 79, 879–888 (2022).

3. Müller, V. I. et al. Altered Brain Activity in Unipolar Depression Revisited: Meta-analyses of Neuroimaging Studies. JAMA Psychiatry 74, 47–55 (2017).

4. Gray, J. P., Müller, V. I., Eickhoff, S. B. & Fox, P. T. Multimodal Abnormalities of Brain Structure and Function in Major Depressive Disorder: A Meta-Analysis of Neuroimaging Studies. Am. J. Psychiatry appiajp201919050560 (2020) doi:10.1176/appi.ajp.2019.19050560.

5. Marek, S. et al. Reproducible brain-wide association studies require thousands of individuals. Nature 603, 654–660 (2022).

6. Schmaal, L. et al. Subcortical brain alterations in major depressive disorder: findings from the ENIGMA Major Depressive Disorder working group. Mol. Psychiatry 21, 806–812 (2016).

7. Schmaal, L. et al. Cortical abnormalities in adults and adolescents with major depression based on brain scans from 20 cohorts worldwide in the ENIGMA Major Depressive Disorder Working Group. Mol. Psychiatry 22, 900–909 (2017).

8. Kendler, K. S., Neale, M. C., Kessler, R. C., Heath, A. C. & Eaves, L. J. A longitudinal twin study of personality and major depression in women. Arch. Gen. Psychiatry 50, 853–862 (1993).

9. Malouff, J. M., Thorsteinsson, E. B. & Schutte, N. S. The relationship between the five-factor model of personality and symptoms of clinical disorders: A meta-analysis. J. Psychopathol. Behav. Assess. 27, 101–114 (2005).

10. Kaiser, R. H., Andrews-Hanna, J. R., Wager, T. D. & Pizzagalli, D. A. Large-Scale Network Dysfunction in Major Depressive Disorder: A Meta-analysis of Resting-State Functional Connectivity. JAMA Psychiatry 72, 603–611 (2015).

11. Koolschijn, P. C. M. P., van Haren, N. E. M., Lensvelt-Mulders, G. J. L. M., Hulshoff Pol, H. E. & Kahn, R. S. Brain volume abnormalities in major depressive disorder: a meta-analysis of magnetic resonance imaging studies. Hum. Brain Mapp. 30, 3719–3735 (2009).

12. Gebre, R. K. et al. Cross-scanner harmonization methods for structural MRI may need further work: A comparison study. Neuroimage 269, 119912 (2023).

13. Casey, B. J. et al. The Adolescent Brain Cognitive Development (ABCD) study: Imaging acquisition across 21 sites. Dev. Cogn. Neurosci. 32, 43–54 (2018).

14. Littlejohns, T. J. et al. The UK Biobank imaging enhancement of 100,000 participants: rationale, data collection, management and future directions. Nat. Commun. 11, 2624 (2020).

15. Glasser, M. F. et al. The Human Connectome Project’s neuroimaging approach. Nat. Neurosci. 19, 1175–1187 (2016).

16. Somerville, L. H. et al. The Lifespan Human Connectome Project in Development: A large-scale study of brain connectivity development in 5-21 year olds. Neuroimage 183, 456–468 (2018).

17. Bookheimer, S. Y. et al. The Lifespan Human Connectome Project in Aging: An overview. Neuroimage 185, 335–348 (2019).

18. Seok, D. et al. Dimensional connectomics of anxious misery, a human connectome study related to human disease: Overview of protocol and data quality. Neuroimage Clin 28, 102489 (2020).

19. Hamilton, M. Development of a rating scale for primary depressive illness. Br. J. Soc. Clin. Psychol. 6, 278–296 (1967).

20. Pilkonis, P. A. et al. Assessment of self-reported negative affect in the NIH Toolbox. Psychiatry Res. 206, 88–97 (2013).

21. Achenbach, T. M. & Edelbrock, C. Child behavior checklist. Burlington (vt) (1991).

22. Dutt, R. K. et al. Mental health in the UK Biobank: A roadmap to self-report measures and neuroimaging correlates. Hum. Brain Mapp. 43, 816–832 (2022).

23. Eysenck, H. J. & Eysenck, S. B. G. Eysenck Personality Questionnaire Manual. (Educational and Industrial Testing Service, San Diego, CA, 1975).

24. Settles, R. E. et al. Negative urgency: a personality predictor of externalizing behavior characterized by neuroticism, low conscientiousness, and disagreeableness. J. Abnorm. Psychol. 121, 160–172 (2012).

25. Klein, A. & Tourville, J. 101 labeled brain images and a consistent human cortical labeling protocol. Front. Neurosci. 6, 171 (2012).

26. Fischl, B. et al. Whole brain segmentation. Neuron 33, 341–355 (2002).

27. Schaefer, A. et al. Local-Global Parcellation of the Human Cerebral Cortex from Intrinsic Functional Connectivity MRI. Cereb. Cortex 28, 3095–3114 (2018).

28. Bayer, J. M. M. et al. Dissecting heterogeneity in cortical thickness abnormalities in major depressive disorder: a large-scale ENIGMA MDD normative modelling study. bioRxivorg 2025.03.17.643677 (2025) doi:10.1101/2025.03.17.643677.

29. Zheng, R., Zhang, Y., Yang, Z., Han, S. & Cheng, J. Reduced brain gray matter volume in patients with first-episode major depressive disorder: A quantitative meta-analysis. Front. Psychiatry 12, 671348 (2021).

30. Peng, W., Chen, Z., Yin, L., Jia, Z. & Gong, Q. Essential brain structural alterations in major depressive disorder: A voxel-wise meta-analysis on first episode, medication-naive patients. J. Affect. Disord. 199, 114–123 (2016).

31. Kandilarova, S., Stoyanov, D., Sirakov, N., Maes, M. & Specht, K. Reduced grey matter volume in frontal and temporal areas in depression: contributions from voxel-based morphometry study. Acta Neuropsychiatr. 31, 252–257 (2019).

32. Goodkind, M. et al. Identification of a common neurobiological substrate for mental illness. JAMA Psychiatry 72, 305–315 (2015).

33. Zhou, H.-X. et al. Rumination and the default mode network: Meta-analysis of brain imaging studies and implications for depression. Neuroimage 206, 116287 (2020).

34. Hamilton, J. P., Farmer, M., Fogelman, P. & Gotlib, I. H. Depressive Rumination, the Default-Mode Network, and the Dark Matter of Clinical Neuroscience. Biol. Psychiatry 78, 224–230 (2015).

35. Gusnard, D. A., Akbudak, E., Shulman, G. L. & Raichle, M. E. Medial prefrontal cortex and self-referential mental activity: relation to a default mode of brain function. Proc. Natl. Acad. Sci. U. S. A. 98, 4259–4264 (2001).

36. Williams, L. M. Precision psychiatry: a neural circuit taxonomy for depression and anxiety. Lancet Psychiatry 3, 472–480 (2016).

37. Hannon, K. et al. Parsing clinical and neurobiological sources of heterogeneity in depression. Biol. Psychiatry (2025) doi:10.1016/j.biopsych.2025.04.025.

38. Hannon, K. et al. Comparing data-driven subtypes of depression informed by clinical and neuroimaging data: A Registered Report. Biol. Psychiatry Glob. Open Sci. 100473 (2025) doi:10.1016/j.bpsgos.2025.100473.

39. Tozzi, L. et al. Reduced functional connectivity of default mode network subsystems in depression: Meta-analytic evidence and relationship with trait rumination. Neuroimage Clin 30, 102570 (2021).

40. Cremers, H. R., Wager, T. D. & Yarkoni, T. The relation between statistical power and inference in fMRI. PLoS One 12, e0184923 (2017).

41. Yeo, B. T. T. et al. The organization of the human cerebral cortex estimated by intrinsic functional connectivity. J. Neurophysiol. 106, 1125–1165 (2011).

42. Videbech, P. & Ravnkilde, B. Hippocampal volume and depression: a meta-analysis of MRI studies. Am. J. Psychiatry 161, 1957–1966 (2004).

43. Belov, V. et al. Multi-site benchmark classification of major depressive disorder using machine learning on cortical and subcortical measures. Sci. Rep. 14, 1084 (2024).

44. Sheline, Y. I., Sanghavi, M., Mintun, M. A. & Gado, M. H. Depression duration but not age predicts hippocampal volume loss in medically healthy women with recurrent major depression. J. Neurosci. 19, 5034–5043 (1999).

45. Sheline, Y. I., Liston, C. & McEwen, B. S. Parsing the Hippocampus in Depression: Chronic Stress, Hippocampal Volume, and Major Depressive Disorder. Biol. Psychiatry 85, 436–438 (2019).

46. Zhang, W., Dutt, R., Lew, D., Barch, D. M. & Bijsterbosch, J. D. Higher amplitudes of visual networks are associated with trait but not state-depression. bioRxiv 2024.03.25.584801 (2024) doi:10.1101/2024.03.25.584801.

47. Wu, F., Lu, Q., Kong, Y. & Zhang, Z. A Comprehensive Overview of the Role of Visual Cortex Malfunction in Depressive Disorders: Opportunities and Challenges. Neurosci. Bull. 1–13 (2023) doi:10.1007/s12264-023-01052-7.

48. Camacho, M. C. et al. Large-scale encoding of emotion concepts becomes increasingly similar between individuals from childhood to adolescence. Nat. Neurosci. 26, 1256–1266 (2023).

49. Javaheripour, N. et al. Altered resting-state functional connectome in major depressive disorder: a mega-analysis from the PsyMRI consortium. Transl. Psychiatry 11, 511 (2021).

50. Kebets, V. et al. Somatosensory-Motor Dysconnectivity Spans Multiple Transdiagnostic Dimensions of Psychopathology. Biol. Psychiatry (2019) doi:10.1016/j.biopsych.2019.06.013.

51. Kent, J. S. et al. Exploring the relationship of transdiagnostic mood and psychosis symptom domains with motor dysfunction. Neuropsychobiology 79, 301–312 (2020).

52. Fritze, S. et al. Deciphering the interplay between psychopathological symptoms, sensorimotor, cognitive and global functioning: a transdiagnostic network analysis. Eur. Arch. Psychiatry Clin. Neurosci. 274, 1625–1637 (2024).

53. Huang, C.-C. et al. Transdiagnostic and Illness-Specific Functional Dysconnectivity Across Schizophrenia, Bipolar Disorder, and Major Depressive Disorder. Biol Psychiatry Cogn Neurosci Neuroimaging 5, 542–553 (2020).

54. Cuijpers, P., Li, J., Hofmann, S. G. & Andersson, G. Self-reported versus clinician-rated symptoms of depression as outcome measures in psychotherapy research on depression: a meta-analysis. Clin. Psychol. Rev. 30, 768–778 (2010).

55. Miguel, C. et al. Self-reports vs clinician ratings of efficacies of psychotherapies for depression: a meta-analysis of randomized trials. Epidemiol. Psychiatr. Sci. 34, e15 (2025).

56. Milak, M. S. et al. Regional brain metabolic correlates of self-reported depression severity contrasted with clinician ratings. J. Affect. Disord. 126, 113–124 (2010).

57. Kawakami, S. et al. Frontal pole-precuneus connectivity is associated with a discrepancy between self-rated and observer-rated depression severity in mood disorders: a resting-state functional magnetic resonance imaging study. Cereb. Cortex 34, bhae284 (2024).

