## Supplementary material for "The neuroimaging correlates of depression established across six large-scale datasets": Online Methods

### Datasets

This study leverages six different datasets, namely the Adolescent Brain Cognitive Development (ABCD) study[1], The UK Biobank (UKB) study[2], the Human Connectome Project (HCP) Young Adult study[3], the HCP Developmental study[4], the HCP Aging study[5], and the Dimensional Connectomics of Anxious Misery study[6]. Subjects with complete and biologically plausible imaging data were included (e.g after removing subjects with any imaging measures equal or greater than five standard deviations away from the dataset mean). In the UKB, an additional set of inclusion criteria was adopted to identify a subset of participants. Specifically, participants were included if they met one or more of three clinical depression criteria: probable MDD status, one or more reported episodes of depression, and/or ICD10 label F31 and F32. The rationale for this UKB subselection was twofold: i) it reduced the storage and computational demands given that the full UKB is the largest sample and ii) it focuses on a more clinically relevant subset of UKB participants. The demographic information for each of the datasets is reported in Table 1.

| | **ABCD** | **HCP-ANXPE** | **UKB** | **HCP-A** | **HCP-D** | **HCP-YA** |
|---|---|---|---|---|---|---|
| Sample size | 9,312 | 185 | 11,000 | 1,132 | 839 | 949 |
| Age (mean±STD) | 9.93±0.62 | 28.45±7.50 | 62.92± 7.46 | 62.48±16.22 | 12.66±2.87 | 28.75±3.73 |
| Sex (% male) | 54 | 34.41 | 40.52 | 44 | 50 | 46 |

*Table 1: Overview of demographics for each of the six datasets*

### Depression Phenotypes

In each dataset, we identified two depression-related phenotypes to use for all linear regression analyses. The depression phenotypes fall into the categories of predisposition and self-reported severity measures (Table 2). The predisposition measures included Eysenck Neuroticism[7] (UKB), NEO five-factors inventory questionnaire Neuroticism subscale (HCP-YA, HCP-A, ANXPE), and UPPS negative urgency[8] (HCP-D and ABCD). The severity measures included Hamilton Depression Rating Scale[9] (ANXPE), NIH Toolbox Sadness[10] (HCP-YA, HCP-A), Child Behavior Checklist (CBCL) Depression subscale[11] (HCP-D, ABCD), and Recent Depressive Symptoms (RDS) scale[12] (UKB).

| | | **ABCD** | **HCP-ANXPE** | **UKB** | **HCP-A** | **HCP-D** | **HCP-YA** |
|---|---|---|---|---|---|---|---|
| Predisposition measures | UPPS negative urgency | x | | | | x | |
| | NEO | | x | | x | | x |
| | Eysenck | | | x | | | |
| Severity measures | CBCL | x | | | | x | |
| | HAM-D | | x | | | | |
| | RDS | | | x | | | |

| | | | | | | | |
|---|---|---|---|---|---|---|---|
| | NIH toolbox sadness | | | | x | | x |

*Table 2: Overview of depression phenotypes measures per dataset in the categories of predisposition and severity.*

### Neuroimaging data and preprocessing

For each study, we leverage T1-weighted structural MRI data and resting state functional MRI data. An overview of the neuroimaging data acquisition parameters can be found in Table 3. We leveraged the preprocessed UKB data in volumetric (nifti) format as shared through the UKB showcase, which has undergone the UKB preprocessing pipeline[13]. We leveraged the preprocessed ABCD data in grayordinate (cifti) format that was preprocessed through adapted HCP-style pipelines[14,15]. For all HCP datasets (HCP-YA, HCP-A, HCP-D, HCP-ANXPE), we leveraged the preprocessed data in grayordinate (cifti) format that was preprocessed through the HCP preprocessing pipeline[16] (including MSM-all registration[17,18] and ICA-FIX cleanup[19,20]).

| | | HCP -ANXPE | UKB | HCP-A | HCP-D | HCP-YA | ABCD | | |
|---|---|---|---|---|---|---|---|---|---|
| | | | | | | | ABCD-1 | ABCD-2 | ABCD-3 |
| | Scanner | Siemens Prisma 3T | Siemens Skyra 3T | Siemens Prisma 3T | Siemens Prisma 3T | Customized 3T ‘Connectom’ | Siemens Prisma 3T | Philips Achieva 3T | GE MR750 3T |
| | Head coil | 64 channel (52 used) | 32 channel | 32 channel | 32 channel | 32 channel | 32 channel | 32 channel | 32 channel |
| **T1w** | Resolution (mm) | .8x.8x.8 | 1x1x1 | .8x.8x.8 | .8x.8x.8 | .7x.7x.7 | 1x1x1 | 1x1x1 | 1x1x1 |
| | FoV (mm) | 256x240x167 | 208x256x256 | 256x240x166 | 256x240x166 | 224x224x | 256x256x176 | 256x256x176 | 256x256x176 |
| | TR (ms) | 2400 | 2000 | 2500 | 2500 | 2400 | 2500 | 6.31 | 2500 |
| | TI (ms) | 1000 | 880 | 1000 | 1000 | 1000 | 1060 | 1060 | 1060 |
| | Scan time (minutes) | XXX | 4:54 | 8:22 | 8:22 | 7:40 | 7:12 | 5:38 | 6:09 |
| **fMRI** | TR (ms) | 800 | 735 | 800 | 800 | 720 | 800 | 800 | 800 |
| | TE (ms) | 37 | 39 | 37 | 37 | 33.1 | 30 | 30 | 30 |
| | Multiband factor | 4 | 8 | 8 | 8 | 8 | 6 | 6 | 6 |
| | Resolution (mm) | 2x2x2 | 2.4x2.4x2.4 | 2x2x2 | 2x2x2 | 2x2x2 | 2.4x2.4x2.4 | 90x90x60 | 90x90x60 |
| | FoV (mm) | 208x208x144 | 211.2x211.2x153.6 | 208x180x144 | 208x180x144 | 208x180x144 | 216x216x144 | 216x216x144 | 216x216x144 |
| | Scan time (minutes) | | 6:10 | | | | | | |

*Table 3: Overview of neuroimaging sequences per dataset*

### Imaging Measures

To facilitate meta analyses across studies, we used the same parcellations between datasets where possible. Specifically, for structural imaging measures, we used the Desikan-Killiany-Tourville (DKT) Atlas to extract cortical thickness, area, and volume measures and the automatic subcortical segmentation (ASEG) to extract subcortical brain volume measures. For the functional imaging measures, we used the Schaefer parcellation (Schaefer) at a dimensionality of 300 parcels[21]. Partial correlation between each possible pair of parcels was calculated as the normalized inverse of the covariance matrix followed by Fisher's r-to-z transformation.

### Mapping parcellations to Yeo networks

For each parcel in the DKT and Schaefer atlases, we calculated the proportion of parcel vertices that overlap with each of the seven Yeo networks. Parcels were assigned to the Yeo network with the largest proportion overlap. All ASEG parcels were combined into one 'subcortical' network. Notably, correlation-based imaging features reflect a pair of parcels and can therefore potentially be mapped onto two different Yeo networks.

### Confound variables

Unless otherwise stated, all regression analyses were controlled for age, sex, total intracranial volume ('ICV'), head motion ('HM'), imaging site (relevant for ABCD and UKB only, coded into sites-1 dummy variables), and family group (relevant for ABCD and HCP-YA only, treated as a random effect). For headmotion, we used the mean rfMRI head motion averaged across space and time points for UKB (variable ID 25741), DVARS median for all HCP datasets (HCP-YA, HCP-A, HCP-D, HCP-ANXPE), and average framewise displacement in mm for ABCD.

### Within-dataset regression analysis and meta analyses

Univariate linear mixed-effects regressions (LMER) were conducted separately for each dataset to assess associations between depression phenotypes ('Dep') and imaging measures ('IDP') as shown in the following formula, where betas are effect sizes for fixed effects and $g$ values are random effects:

$$IDP = β\_1*Dep + β\_2*age + β\_3*sex + β\_4*HM + β\_5*ICV + g\_1*site + g\_2*familyID + ε$$

No intersect term was included as all input data were normalized prior to regression analyses. The regression coefficients of interest ($β_1$) and associated error term were subsequently entered into separate meta analyses for each depression phenotype and imaging measure to calculate the meta analytical effect size, meta analytical p-value, and meta analytical confidence interval. False discovery rate correction was applied to the meta analytical p-values within each depression phenotype to control for multiple comparisons across all 45,052 imaging measures

(44,850 functional connectivity, 62 cortical thickness, 62 cortical surface area, and 78 gray matter volume).

### Statistical comparisons

Meta analytic effect sizes were entered into several analyses of variance (ANOVAs) for further comparison. Absolute values of the meta-analytical effect sizes were used as the inputs for all ANOVAs to avoid negative and positive effects canceling out. Specifically, one ANOVA was performed to statistically compare structural versus functional imaging measures and two separate ANOVAs (for two imaging metric types) were used to assess the spatial distribution and the difference between depression phenotypes, as described below:

1. A two-way ANOVA with a main effect for imaging metric type (4 levels; gray matter volume, cortical surface area, cortical thickness, functional connectivity), a main effect for depression phenotype (2 levels; severity, predisposition), and the interaction effect (imaging metric type x depression phenotype) was performed to robustly compare between structural and functional imaging measures.
2. Two separate two-way ANOVAs with a main effect for Yeo network (7 or 8 levels; depending on the inclusion/exclusion of subcortical regions), a main effect for depression phenotype (2 levels; severity and predisposition), and the interaction effect (Yeo network x depression phenotype) were performed to assess the role of spatial distribution and depression phenotype. To ensure interpretability, these ANOVAs were performed separately for gray matter volume and cortical surface area. Equivalent analyses were not performed for cortical thickness or functional connectivity because neither of these imaging metric types resulted in substantive significant meta analytical results.

**References**


1. Casey, B. J. *et al.* The Adolescent Brain Cognitive Development (ABCD) study: Imaging acquisition across 21 sites. *Dev. Cogn. Neurosci.* **32**, 43–54 (2018).
2. Littlejohns, T. J. *et al.* The UK Biobank imaging enhancement of 100,000 participants: rationale, data collection, management and future directions. *Nat. Commun.* **11**, 2624 (2020).
3. Glasser, M. F. *et al.* The Human Connectome Project's neuroimaging approach. *Nat. Neurosci.* **19**, 1175–1187 (2016).
4. Somerville, L. H. *et al.* The Lifespan Human Connectome Project in Development: A large-scale study of brain connectivity development in 5-21 year olds. *Neuroimage* **183**,

456–468 (2018).

5. Bookheimer, S. Y. *et al.* The Lifespan Human Connectome Project in Aging: An overview. *Neuroimage* **185**, 335–348 (2019).

6. Seok, D. *et al.* Dimensional connectomics of anxious misery, a human connectome study related to human disease: Overview of protocol and data quality. *Neuroimage Clin* **28**, 102489 (2020).

7. Eysenck, H. J. & Eysenck, S. B. G. *Eysenck Personality Questionnaire Manual*. (Educational and Industrial Testing Service, San Diego, CA, 1975).

8. Settles, R. E. *et al.* Negative urgency: a personality predictor of externalizing behavior characterized by neuroticism, low conscientiousness, and disagreeableness. *J. Abnorm. Psychol.* **121**, 160–172 (2012).

9. Hamilton, M. Development of a rating scale for primary depressive illness. *Br. J. Soc. Clin. Psychol.* **6**, 278–296 (1967).

10. Pilkonis, P. A. *et al.* Assessment of self-reported negative affect in the NIH Toolbox. *Psychiatry Res.* **206**, 88–97 (2013).

11. Achenbach, T. M. & Edelbrock, C. Child behavior checklist. *Burlington (vt)* (1991).

12. Dutt, R. K. *et al.* Mental health in the UK Biobank: A roadmap to self-report measures and neuroimaging correlates. *Hum. Brain Mapp.* **43**, 816–832 (2022).

13. Alfaro-Almagro, F. *et al.* Image processing and Quality Control for the first 10,000 brain imaging datasets from UK Biobank. *Neuroimage* **166**, 400–424 (2018).

14. Goncalves, M. *et al.* FMRIPrep Lifespan: Extending A robust pipeline for functional MRI preprocessing to developmental neuroimaging. *bioRxiv* 2025.05.14.654069 (2025) doi:10.1101/2025.05.14.654069.

15. Feczko, E. *et al.* Adolescent brain cognitive development (ABCD) community MRI collection and utilities. *bioRxiv* 2021.07.09.451638 (2021) doi:10.1101/2021.07.09.451638.

16. Glasser, M. F. *et al.* The minimal preprocessing pipelines for the Human Connectome

Project. *Neuroimage* **80**, 105–124 (2013).

17. Robinson, E. C. *et al.* Multimodal surface matching with higher-order smoothness constraints. *Neuroimage* **167**, 453–465 (2018).
18. Robinson, E. C. *et al.* MSM: a new flexible framework for Multimodal Surface Matching. *Neuroimage* **100**, 414–426 (2014).
19. Salimi-Khorshidi, G. *et al.* Automatic denoising of functional MRI data: combining independent component analysis and hierarchical fusion of classifiers. *Neuroimage* **90**, 449–468 (2014).
20. Griffanti, L. *et al.* ICA-based artefact removal and accelerated fMRI acquisition for improved resting state network imaging. *Neuroimage* **95**, 232–247 (2014).
21. Schaefer, A. *et al.* Local-Global Parcellation of the Human Cerebral Cortex from Intrinsic Functional Connectivity MRI. *Cereb. Cortex* **28**, 3095–3114 (2018).
